# Multiple clones of colistin-resistant *Salmonella enterica* carrying *mcr-1* plasmids in meat products and patients in Northern Thailand

**DOI:** 10.1101/2020.12.08.415869

**Authors:** Prapas Patchanee, Nipa Chokesajjawatee, Pannita Santiyanont, Phongsakorn Chuammitri, Manu Deeudom, William Monteith, Samuel K. Sheppard, Ben Pascoe, Teerarat Prasertsee

## Abstract

*Salmonella* spp. is an important foodborne pathogen associated with consumption of contaminated food, especially livestock products. Antimicrobial resistance (AMR) in *Salmonella* has been reported globally and increasing AMR in food production is a major public health issue worldwide. The objective of this study was to describe the genetic relatedness among *Salmonella enterica* isolates, which displayed identical DNA fingerprint profiles. Ten *S. enterica* isolates were selected from meat and human cases with an identical rep-PCR profile of serovars Rissen (n=4), Weltevreden (n=4), and Stanley (n=2). We used long-read whole genome sequencing (WGS) on the MinION sequencing platform to type isolates and investigate *in silico* the presence of specific AMR genes. Antimicrobial susceptibility testing was tested by disk diffusion and gradient diffusion method to corroborate the AMR phenotype. Multidrug resistance and resistance to more than one antimicrobial agent were observed in eight and nine isolates, respectively. Resistance to colistin with an accompanying *mcr*-1 gene was observed among the *Salmonella* isolates. The analysis of core genome and whole genome MLST revealed that the *Salmonella* from meat and human salmonellosis were closely genetic related. Hence, it could be concluded that meat is one of the important sources for *Salmonella* infection in human.

**Highlights:**

- Colistin resistance detected in 2 clones from 2 different *Salmonella enterica* serovars (Rissen and Weltevreden) with accompanying plasmid-borne *mcr*-1 gene from the food production chain and human clinical salmonellosis.
- High prevalence of multidrug resistant isolates and resistance to more than one antimicrobial agent.
- MinION has potential for mobile, rapid and accurate application in veterinary genomic epidemiology studies.

## 1. Introduction

Non-typhoidal *Salmonella* (NTS) is a major cause of foodborne-related disease globally. In farm animals, NTS infection is found in a high prevalence in South-East Asia (Carrique-Mas and Bryant, 2013). Infected farm animals are the major reservoir of *Salmonella* infection and are considered the original source of slaughterhouse contamination (Trongjit et al., 2017). High density of livestock production on farms, improper hygiene practice in abattoirs and retail stores, and limited cold chain protection during meat distribution can support pathogen growth at each stage of the production chain (Heredia and García, 2018). NTS can transmit to humans by consumption of contaminated food, including consumption of raw or undercooked meat, which is the most important risk factor associated with *Salmonella* infection in humans (Padungtod and Kaneene, 2006). Annually, over 90 million people fall ill and 155,000 deaths are caused by NTS infection (Heredia and García, 2018). Salmonellosis caused by non-invasive NTS serovars are usually self-limiting, but severe infection can occur in young children, the elderly and immunocompromised individuals. Complications with infection such as bacteraemia occur most often in high-risk groups of patients, thus antimicrobial agents will often be used for medical treatment (Prasertsee et al., 2019).

Antimicrobial resistance (AMR) in food-borne pathogens has been reported with high frequency and poses a significant public health threat (Prestinaci et al., 2015). Overuse and misuse of antimicrobial agents in livestock production can lead to the development of antimicrobial resistance in bacteria (Trongjit et al., 2017). Specifically, colistin is a polymyxin (Srinivas and Rivard, 2017) that is used in both human and veterinary medicine against Gram-negative bacterial infections (Magiorakos et al., 2012). Although the mechanism of resistance is not fully characterised, colistin is able to disrupt lipopolysaccharides and phospholipids in the outer membrane of Gram-negative bacteria, causing cell death (Biswas et al., 2012). Among other antimicrobial agents in veterinary medicine, in addition to treatment colistin is often used prophylactically in food-producing animals for growth promotion (Catry et al., 2015). This has had global implications on the emergence of AMR in food production this century (Biswas et al., 2012; Exner et al., 2017; Iwu et al., 2016).

Prevention and control of *Salmonella* infection requires comprehensive data collection for effective surveillance systems. Sequencing of bacterial whole genomes provides a means for detection of virulence genes, AMR genes and plasmid replicons within a single run. In this study, we use long-read sequencing technology to characterize selected MDR *Salmonella enterica* clones from meat and human cases. Oxford Nanopore is portable and offers real-time sequencing that can be performed with minimal specialist equipment (Gardy et al., 2015), and allows us to identify plasmid-borne antibiotic resistance genes. We aimed to assess the diversity of plasmids carrying *mcr-1*, which mediates colistin resistance, and consider the threat to public health posed by the presence of these plasmids in the local food production chain.

## 2. Materials and Methods

### 2.1. Bacterial strains

Ten *Salmonella* isolates were selected from three clones with previously published identical rep-PCR profiles (Prasertsee et al., 2019). Selected *Salmonella* isolates comprised of *S*. Rissen (n=4), *S*. Weltevreden (n=4), and *S*. Stanley (n=2) clones which were isolated from meat (chicken meat, pork, and beef) and human cases in Maharaj Nakhon Chiang Mai Hospital from January to July, 2017. Ethical approval for collection of *Salmonella* isolated from humans in this study was approved by the Human Research Ethics Committee, Faculty of Medicine, Chiang Mai University (Study code: NONE-2560-04782/Research ID: 4782). Culturing was permitted by Institutional Biosafety Committee-Chiang Mai University (approval No.CMUIBC A-0560001).

### 2.2. Antimicrobial susceptibility testing and minimum inhibitory concentrations

Antimicrobial susceptibility testing and minimum inhibitory concentrations (MICs) of all *Salmonella* isolates were performed at the Central Laboratory, Faculty of Veterinary Medicine, Chiang Mai University. Disc diffusion method and MICs were tested for 10 antimicrobial agents as follows: ampicillin (AMP, 10 μg), amoxycillin/clavulanate (AMC, 20/10 μg), chloramphenicol (CHL, 30 μg), ciprofloxacin (CIP, 5 μg), nalidixic acid (NAL, 30 μg), norfloxacin (NOR, 10 μg), streptomycin (STR 10 μg), sulfisoxazole (SX, 250 μg), tetracycline (TET, 30 μg), and trimethoprim/sulfamethoxazole (SXT, 1.25/23.75 μg). Inhibition zones were measured according to the guidelines of the Clinical and Laboratory Standards Institute (CLSI, 2017). *Escherichia coli* ATCC 25922 was used as an internal quality control.

European Committee on Antimicrobial Susceptibility Testing (EUCAST, 2016) guidelines were used to determine susceptibility to colistin (COL) by the gradient diffusion method. Results were interpreted using the Annex 1: EUCAST clinical breakpoints and epidemiological cut-off values for the priority list of antimicrobials to be tested for *Salmonella* spp. (MIC dilution ≤ 2 mg/L was recorded as sensitive, while MICs > 2 mg/L were considered resistant).

### 2.3. DNA extraction and Whole genome sequencing

*Salmonella* isolates were cultured overnight in nutrient broth (NB; Merck, Darmstadt, Germany at 37°C. DNA was extracted using the Wizard Genomic DNA Purification Kit, according to manufacturer’s protocols (Promega, Madison, WI, USA). DNA purity and concentration were quantified using a NanoDrop spectrophotometer (Thermo Fisher Scientific, Waltham, MA, USA). DNA concentrations ranged from 61.1 to 655.3 ng/ul.

Sequencing libraries were prepared using the MinION Genomic DNA Sequencing Kit (Oxford Nanopore Technologies, Oxford, UK) with the Rapid Barcoding Kit (SQK-RBK004) to barcode individual samples according to the manufacturer’s instructions. The MinION flow cell (R9 flow cell chemistry) was inserted into the MinION device and connected to the computer (Windows 10; USB 3.0; SSD; i7 processor) via USB 3.0. The MinKNOW v1.15.4 software was run to sequence the *Salmonella* genome for 48 hr. Albacore v2.0.1 was used to call bases and convert FAST5 (Nanopore raw reads) to FASTQ format. Raw reads shorter than 500 base pairs or quality scores less than 7 were filtered with Nanofilt v.1.0.5. Porechop v.0.2.3 was used to remove the internal barcode adapter sequences and genomes were assembled using Unicycler v0.4.2. The resulting FASTA files of each *Salmonella* isolate was scanned to identify AMR genes and determine cg/wgMLST profiles.

### 2.4. *In silico* antimicrobial resistance gene detection

Ten assembled *Salmonella* genomes were probed for the presence of AMR genes using the ResFinder 3.1 database with the parameters: 90% identification threshold and 40% minimum length. The AMR genes in this database included aminoglycoside (*aadA1, aadA2, aph3, aph6, and strA*), beta-lactam (*bla*_TEM-1B_), quinolone (*qnrS1*), macrolide (*mph(A)* and *mef(B)*), phenicol (*cmlA1, cml, and floR*), polymyxin (*mcr-1*), sulfonamide (*sul1, sul2*, and *sul3*, tetracycline (*tet(A)* and *tet(M)*, and trimethoprim (*dfrA12*) (https://cge.cbs.dtu.dk.services/ResFinder/) (Zankari et al., 2012).

### 2.5. Whole genome and core genome MLST (wgMLST and cgMLST) analysis

The wgMLST and cgMLST profiles were characterised using WGS tools plugin and assembly-based allele calling algorithms in the BioNumerics v.7.6.3 software (Applied Maths, Sint-Martens-Latem, Belgium). The wgMLST schema included 15,874 loci generated from previously published *S. enterica* reference genomes. The cgMLST analysis included 1,509 loci which represented more than 80% of the genes in more than 95% of our *Salmonella* isolates (Vincent et al., 2018). Within *Salmonella* isolates, 1,916 loci represented the loci that are found in at least one isolate with more than 80% homology. The dendrograms of cgMLST and wgMLST were calculated by categoricl differences as a similarity coefficient and using the UPGMA clustering method.

### 2.6. *In silico* plasmid typing

The MOB-suite software was used to type the plasmid sequences (Robertson and Nash, 2018). The MOB-suite software includes the MOB-typer tool, which provides replicon typing for categorizing plasmids based on the sequences responsible for plasmids replication. Furthermore, the MOB-typer provides a prediction of plasmid transmissibility. Based on the presence of relaxase, mate-pair formation and *oriT* sequences, plasmid transmissibility was predicted as either “conjugative” (i.e. sequences including both a relaxase and a mate-pair formation marker), “mobilizable” (i.e. sequences including only a relaxase or an *oriT*) or “non-mobilizable” (i.e. both relaxase and an *oriT* are absent from the sequence). Additionally, MOB-suite provides a scalable nomenclature for plasmid identification by estimating genomic distances using Mash min-hashing.

## 3. Results

### 3.1. Whole genome and core genome MLST (wgMLST and cgMLST) analysis

A total of ten *Salmonella* isolates from different meat types and salmonellosis patients were sequenced by MinION. The cg and wgMLST schemes were analysed by 1,509 and 1,916 loci shared within all *Salmonella* isolates. **Figure 1** demonstrated the phylogenetic tree of cg and wgMLST analysis. Each of *Salmonella* serotype was clustered separately. Identical clones (100% similarity) were not observed in the cg and wgMLST analyses. Based on cgMLST analysis, the *Salmonella* isolates were closely related within each serovar, sharing more than 98% similarity. These results were the same as wgMLST which showed a genetic difference of less than 2% within each group of serovar.

**Figure 1.**
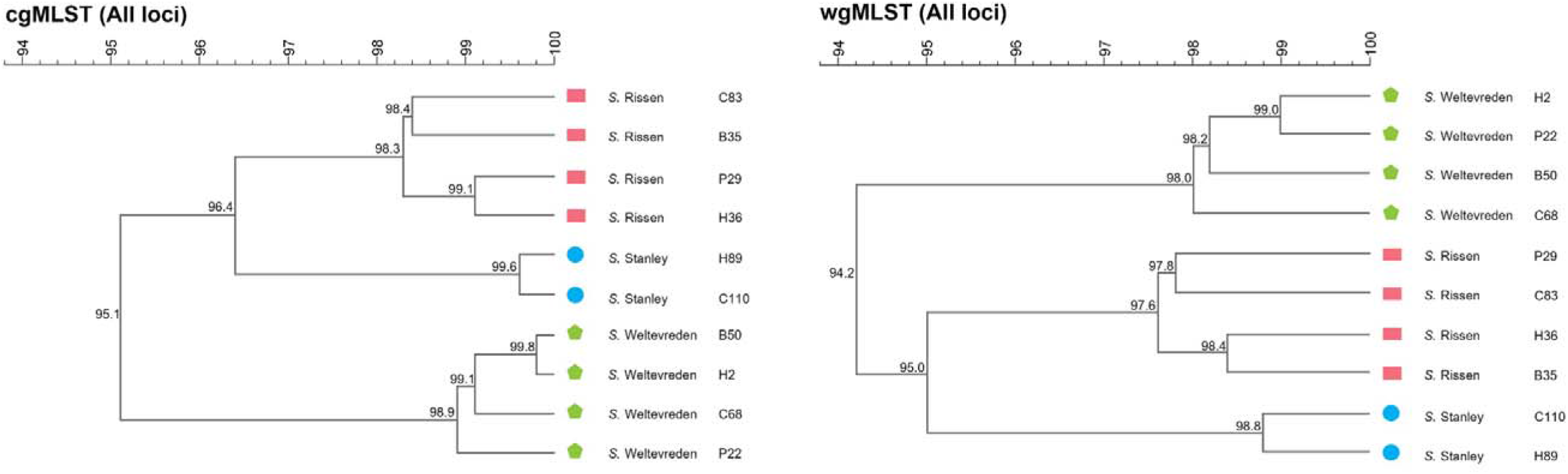
The analysis of cgMLST and wgMLST of ten *Salmonella* isolates from various type of meat and human cases in this study. The tree was generated by using the core genome and whole genome MLST schemes in BioNumerics software. The numbers on each branch illustrate the similarity index. Colors and symbols indicate the *Salmonella* serotype: *S*. Rissen (red rectangle); *S*. Stanley (blue circle); and *S*. Weltevreden (green pentagon).

### 3.2. Antimicrobial susceptibility testing and minimum inhibitory concentrations (MICs)

Antimicrobial susceptibility was tested by the disc diffusion method and the results were determined according to CLSI guidelines, 2017. MICs for eight of our ten *Salmonella* isolates were higher than the CLSI resistance breakpoint for ampicillin, while all isolates had an MIC lower than 8/4 μg/ml (regarded as susceptible by CLSI, 2017) for amoxicillin-clavulanic acid (**Table 1**). One isolate was resistant to quinolones, as demonstrated by a ciprofloxacin MIC above 1 μg/ml, two isolates were resistant to nalidixic acid (>32 μg/ml), and three isolates showed resistance to norfloxacin (>16 μg/ml) (**Table 1**). Additionally, the MICs towards tetracycline and chloramphenicol were higher than the CLSI resistance breakpoints in more than 50% of isolates (6/10 and 5/10, respectively; **Table 1**).

**Table 1:** MIC profile of ten *Salmonella* isolates

| Antimicrobial Agents | MIC (µg/ml) |  |  |  |  |  |  |  |  |  |
| --- | --- | --- | --- | --- | --- | --- | --- | --- | --- | --- |
|  | Rissen |  |  |  | Weltevreden |  |  |  | Stanley |  |
|  | H36 | B35 | C83 | P29 | H2 | B50 | C68 | P22 | H89 | C110 |
| AMP<sup>1</sup> | >32 | >32 | >32 | >32 | <8 | >32 | <8 | >32 | >32 | >32 |
| AMC<sup>1</sup> | <8/4 | <8/4 | <8/4 | <8/4 | <8/4 | <8/4 | <8/4 | <8/4 | <8/4 | <8/4 |
| CHL<sup>1</sup> | <8 | <8 | <8 | >32 | >32 | <8 | >32 | <8 | >32 | >32 |
| CIP<sup>1</sup> | <0.06 | <0.06 | <0.06 | <0.06 | >1 | <0.06 | <0.06 | <0.06 | <0.06 | <0.06 |
| NAL<sup>1</sup> | >32 | <16 | <16 | <16 | <16 | <16 | <16 | <16 | >32 | <16 |
| NOR<sup>1</sup> | >16 | <4 | <4 | <4 | >16 | <4 | <4 | <4 | >16 | <4 |
| STR<sup>1</sup> | <32 | <32 | <32 | <32 | <32 | >64 | >64 | <32 | >64 | <32 |
| SX<sup>1</sup> | <256 | <256 | <256 | <256 | <256 | <256 | >512 | >512 | <256 | >512 |
| TET<sup>1</sup> | <4 | >16 | <4 | >16 | >16 | >16 | <4 | >16 | >16 | <4 |
| SXT<sup>1</sup> | >4/76 | <2/38 | <2/38 | <2/38 | <2/38 | >4/76 | >4/76 | >4/76 | <2/38 | <2/38 |
| COL<sup>2</sup> | <2 | <2 | <2 | >2 | >2 | <2 | <2 | >2 | <2 | <2 |

<sup>1</sup> Breakpoints established by CLSI, 2017
<sup>2</sup> Clinical breakpoints established by EUCAST, 2016

All three *Salmonella* clones were extensively multidrug resistant (**Table 2**). Resistance to more than one antimicrobial agent was observed in nine samples and multidrug resistance (resistance to at least one agent in three or more antimicrobial categories) was found in eight isolates. In this study, resistance to six antimicrobial agents (AMP, CHL, STR, TET, NOR, NAL) was observed in a Stanley serovar isolate (H89), which was isolated from a human clinical case. Extensive-multidrug resistance was observed in four isolates (H36, H2, B50 and 1’22). which were resistance to five antimicrobial agents. None of the *Salmonella* samples in this study were susceptible to all antimicrobial agents.

**Table 2:** The antimicrobial resistance profiling and resistance genes of the *Salmonella* isolates in this study

| Serotypes | ID | Antimicrobial resistance phenotypes | Antimicrobial resistance genes |  |  |
| --- | --- | --- | --- | --- | --- |
|  |  |  | Chromosome | Plasmid1 | Plasmid2 |
| Rissen | H36 | AMP, SXT, TET, NOR, NAL | *tet(A), sul1, sul3, cmlA1, bla<sub>TEM-207</sub>* | *qnrS1* | - |
|  | B35 | AMP, TET | *aadA1, bla<sub>TEM-148</sub>, bla<sub>TEM-208</sub>, dfrA12* | - | - |
|  | C83 | AMP | - | - | - |
|  | P29 | AMP, CHL, TET, COL | *aac(6')-Iaa* | *cmlA* | *mcr-1* |
| Weltevreden | H2 | CHL, TET, NOR, CIP, COL | - | *mcr-1* | - |
|  | B50 | AMP, STR, SXT, TET, NOR | *aac(6')-Iaa* | *mcr-1* | *dfrA12* |
|  | C68 | CHL, STR, SX, SXT | *aac(6')-Iaa* | *qnrS1, qnrS3* | - |
|  | P22 | AMP, SX, SXT, TET, COL | - | *mcr-1* | - |
| Stanley | H89 | AMP, CHL, STR, TET, NOR, NAL | *bla<sub>TEM-128</sub>, bla<sub>TEM-198</sub>, bla<sub>TEM-1B</sub>, tet(M)* | *qnrS1, qnrS7* | - |
|  | C110 | AMP, CHL, SX | *aac(6')-Iaa* | *aadA1, aadA2, cmlA1, dfrA12, sul3* | - |

Fluoroquinolones are recommended as the first line therapeutic drugs for NTS infection. In this study, resistance to fluoroquinolones was also observed in all three serovars. Resistance to quinolones (ciprofloxacin, nalidixic acid, and norfloxacin) was present in *Salmonella* isolated from human cases (H36, H2, and H89) and chicken meat (C83) (**Table 2**). Four isolates showed resistance to norfloxacin, two were resistant to nalidixic acid, and one Weltevreden serovar isolate (H2) was resistant to ciprofloxacin (second-generation quinolones). In addition, resistance to colistin (MICs >2 μg/ml) was found in serovars Rissen (P29) and Weltevreden (H2 and P22).

Long-read sequencing provided us with ten closed, circular genomes and in the Rissen and Stanley isolates additional plasmid DNA was also sequenced, including two plasmids each for isolates P29 and B50 (**Table 3**). The most common antibiotic resistance gene we identified was the *bla*_TEM-1B_ gene, which regulates beta-lactam resistance and was identified in nine out of ten isolates. Five *Salmonella* isolates contained the tetracycline resistance gene, *tetA*. Four out of ten isolates contained the chloramphenicol exporter *cmlA1*, aminoglycoside resistance gene *aadA2*, integron-encoded dihydrofolate reductase *drfA12* and the colistin resistance gene *mcr-1*. In addition, three *Salmonella* isolates (H36, P22, and H89) carried the *qnrS1* gene, which is involved in fluoroquinolone resistance.

**Table 3:**
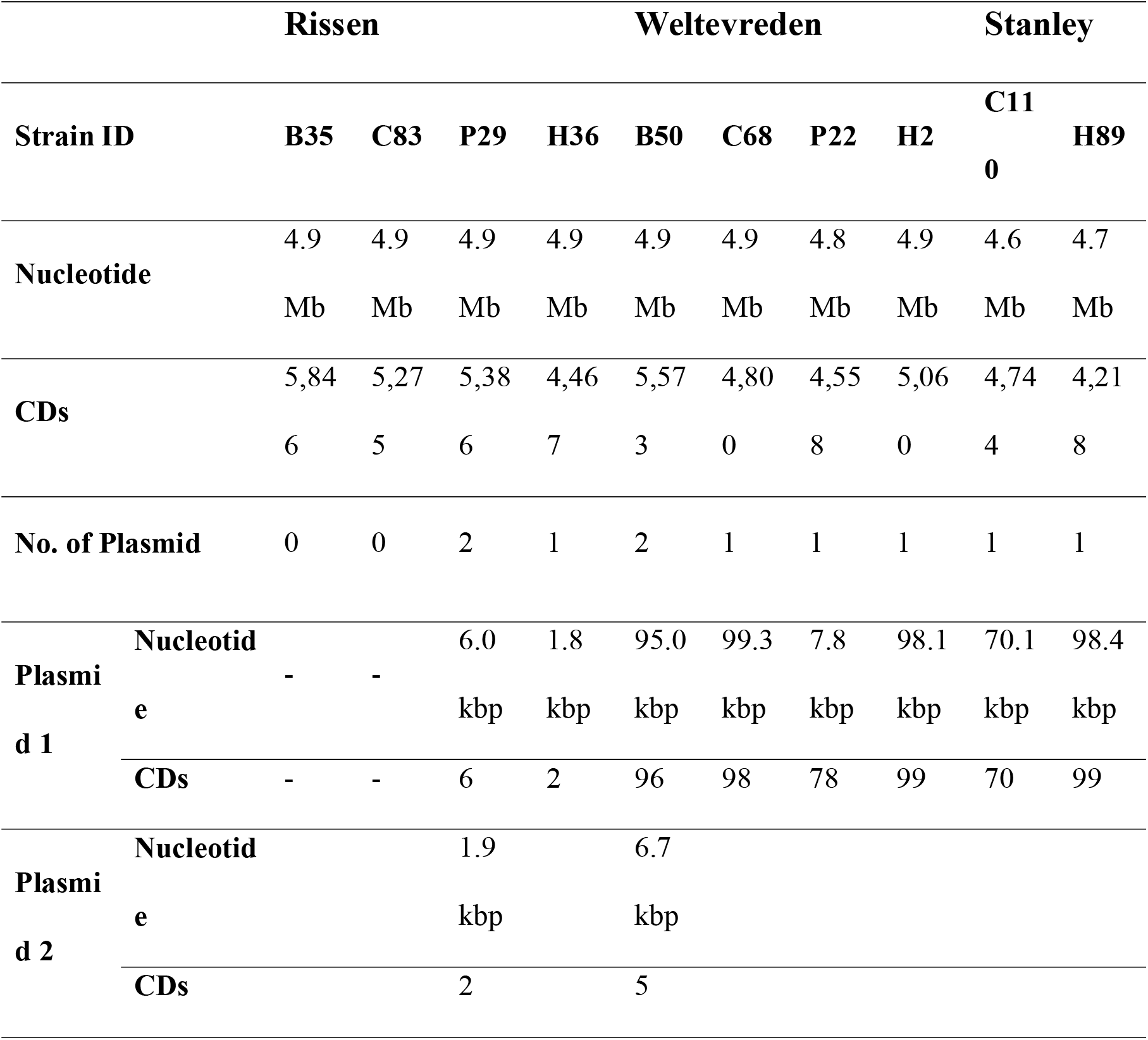
Summary of the whole genome sequencing data and their plasmid insight

### 3.3. Detection and potential transmission of multiple *mcr-1* plasmids in meat production and human salmonellosis

Presence of the colistin resistance gene *mcr-1* does not always result in phenotypic resistance, sometimes the gene is not expressed or requires additional elements to confer resistance (Ahmed et al., 2019). Colistin resistance genes were reported in serovar Rissen and Weltevreden and two of the *Salmonella* isolates from pork (P29, P22) and one human salmonellosis case (H2) demonstrated corresponding phenotypic resistance. However, one of the four isolates (B50) from beef that contained the *mcr-1* gene did not express the colistin resistance phenotype (**Table 2**).

Plasmid sequences from the *Salmonella* isolates P29, H2, B50, and P22 carried the *mcr-1* gene, which can potentially confer colistin resistance. Several different *mcr-1* plasmids have been described containing different incompatibility (Inc) groups and additional antimicrobial resistance determinants. MOB-typer identified 3 distinct *mcr-1* plasmids in our collection. Two (H2 and B50) contain the IncFII replicase and are predicted to be mobilizable given that they contain oriT sequences but lack a mate-pair formation sequence. Plasmid clustering revealed the H2 and B50 *mcr-1* plasmids to both be genetically similar to an IncF-type plasmid (NCBI accession number: LN890519) based on the min-hash clustering method. The P22 plasmid was also predicted to be mobilizable but contained a distinct replicase type compared to the H2 and B50 *mcr-1* plasmids. The P29 plasmid was predicted to be non-mobilizable and MOB-typer failed to identify a replicase site within the sequence (**Table 4**).

**Table 4:**
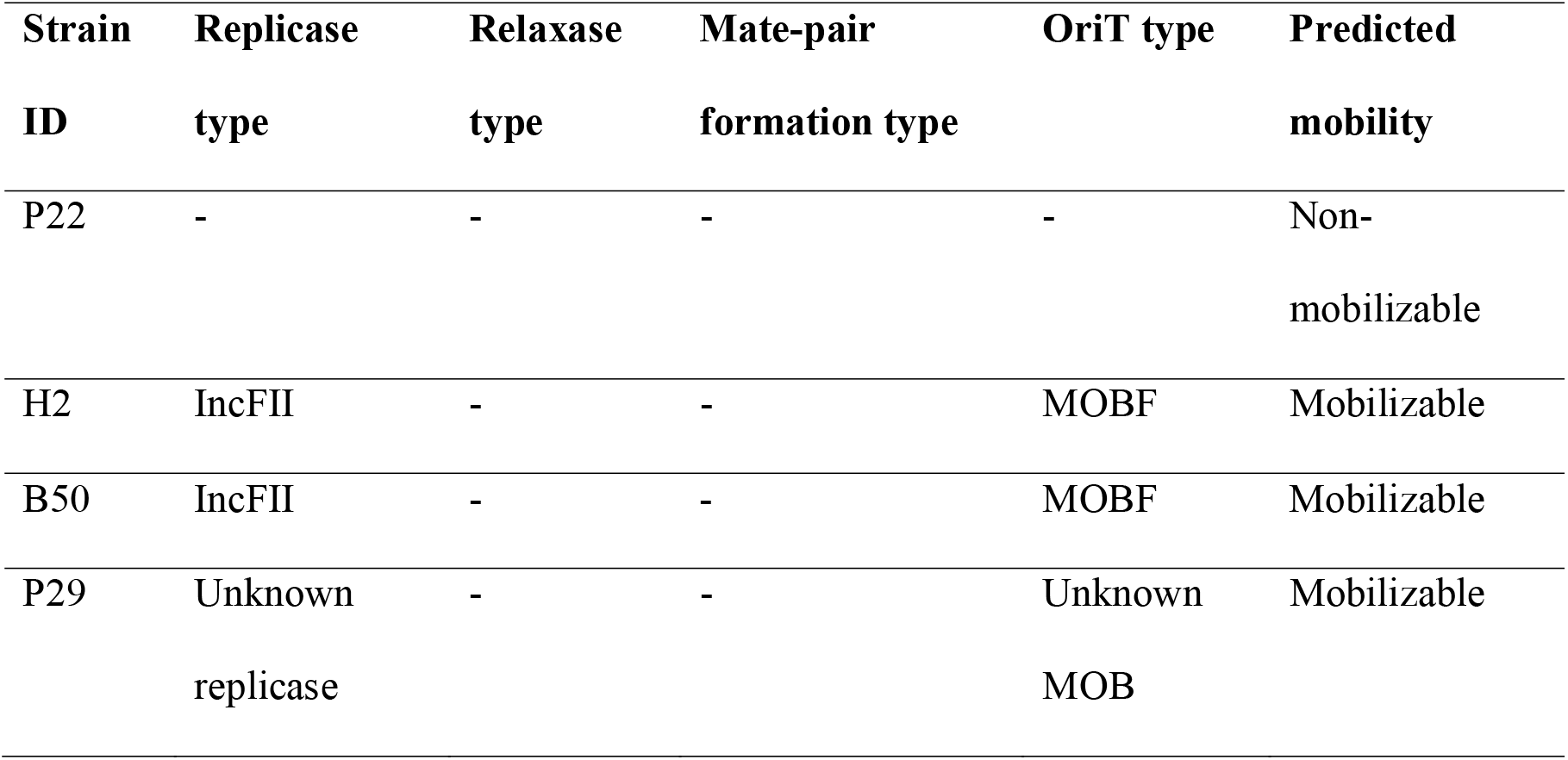
Plasmid typing of the *mcr-1* plasmids

### 3.4. Additional antimicrobial resistant genes

An aminoglycoside resistance gene *(aac(6’)-Iaa)* was frequently detected in the chromosome of four of our *Salmonella* isolates (P29, B50, C68, and C110). Tetracycline (*tet(A), tet(M)*) and beta-lactam (*bla*_TEM-207_, *bla*_TEM-148_, *bla*_TEM-208_, *bla*_TEM-128_, *bla*_TEM-198_, and *bla*_TEM-1B_) resistance genes were also found integrated in the chromosomes of *Salmonella* in this study. Fluoroquinolone resistance genes: *qnrS1, qnrS3, and qnrS7* were found only in the plasmid DNA of the *Salmonella* isolates H36, C68 and H89 (Table 2). Other aminoglycoside resistance genes (*aadA1, aadA2*), chloramphenicol resistance gene (*cmlA1*), sulfonamide and trimethoprim (*sul1, sul3, and dfrA12*) could be detected in both chromosome and plasmid sequences (**Table 2 and 3**).

## 4. Discussion

The dramatic rise of AMR in bacteria could account for up to 10 million associated deaths by 2050, if no action is taken (Balouiri et al., 2016). Transmission between niches (in this case through the food production chain) can complicate effective intervention. In this study, by focusing analyses on individual resistant clones and associated plasmids, we characterize directly the means by which multidrug resistance has been spread between pathogenic *Salmonella* serovars common in the food production chain. Multidrug resistance plasmids were found in high rate (8/10) and occurred in every serotype. the heavy use of related antimicrobials in human and veterinary medicine (Livermore et al., 2007; Schwarz et al., 2001; Teuber, 2001), has raised concerns about how selection for resistance in livestock may lead to AMR in human pathogens. Despite the ban on the use of antibiotics as growth promoters in animals in 2006, quinolones and tetracycline are still available for treatment of livestock all over the world.

Multidrug resistant *Salmonella* have been reported frequently in SE Asia, and in 2017 multidrug resistant *Salmonella* were found to be common in pig and chicken products in Eastern Thailand and Cambodia border provinces (Trongjit et al., 2017). Moreover, a high prevalence of multidrug resistance among *Salmonella* isolates from pork and chicken meat at retail markets has also been observed in Guangdong, China (Zhang et al., 2018). In northeastern Thailand and Laos, as many as 92% (Thailand) and 100% (Laos) of samples from human cases were identified as multidrug resistant (Sinwat et al., 2016). Potential misuse or/and overuse of antimicrobial agents in human and food animals is likely a major contributor to increasing rates of AMR observed this decade (Wang et al., 2019).

Colistin resistance in bacteria isolated from livestock animals, environmental sources and human clinical cases has been widely reported over the world. In this study, *S*. Rissen (P29) and *S*. Weltevreden (H2, P22) were resistant to colistin and carried *mcr-1* gene which regulated the colistin resistance. Several reports have described the presence of *mcr-1* gene in *Enterobacteriaceae* isolated from animals and humans worldwide (Du et al., 2016; Liu et al., 2016; Tuo et al., 2018). The *mcr-1* gene was located on the plasmid such as IncX4 and IncHI2, these two plasmids have been associated with spreading *mcr-1* in *Salmonella* spp. and other bacteria *Enterobacteriaceae* (Campos et al., 2016; Doumith et al., 2016; Skov and Monnet, 2016; Veldman et al., 2016). In this study, we identified 3 distinct plasmids carrying *mcr-1*, two (H2 and B50) contained the IncFII replicase and are predicted to be mobile. Comparison of plasmid sequences revealed the H2 and B50 *mcr-1* plasmids to be genetically similar to an IncF-type plasmid (NCBI accession number: LN890519) previously identified in Southeast Asia (Makendi et al., 2016). The P22 plasmid was also predicted to be mobile but contained a distinct replicase type compared to the H2 and B50 *mcr-1* plasmids. The P29 plasmid was predicted to be non-mobile and MOB-typer failed to identify a replicase site within the sequence (Table 4).

A very high rate of resistance to ampicillin (8/10) and tetracycline (7/10) was observed in this study, similar to previous results from northeastern Thailand, Laos and Vietnam where ampicillin and tetracycline resistance was common in *S. enterica* isolated from food animals and their products (Sinwat et al., 2016; Thai et al., 2012). This finding is also supported by evidence of high rates of ampicillin and tetracycline use for growth promotion, prophylaxis, treatment and infection control in livestock animals (Divek et al., 2018; Nhung et al., 2018).

Chloramphenicol use in livestock production has been prohibited in Thailand, and many other countries (Berendsen et al., 2010). However, resistance to chloramphenicol is still reported in 50% (5/10) of the isolates in this study. The *cmlA* gene, which is associated with chloramphenicol resistance, was detected in four *Salmonella* isolates and three isolates (P29, C68, and H89) demonstrated corresponding phenotypic chloramphenicol resistance The *cmlA* gene is a part of the gene cassette carried by class 1 integrons, which is expressed to provide chloramphenicol resistance. A lack of gene expression may explain why the gene can be detected in one of our isolates without corresponding phenotypic resistance (Chuanchuen and Padungtod, 2009). Prior exposure to antibiotics can increase the selective pressure for carriage and expression of these antimicrobial determinants, which correlates with high rates of antibiotic usage in the region (Prasertsee et al., 2016).

Fluoroquinolones are the first-line antibiotic for treatment of diarrheal disease in both human and domestic animals (Collignon, 2005; Tribble, 2017). Our findings reveal that all *Salmonella* isolated from human cases were resistance to norfloxacin. Additionally, all *S*. Weltevreden isolated from human cases were also phenotypically resistant to ciprofloxacin, the second-generation quinolones group. Isolate B50 was the only isolate from meat which was norfloxacin resistant. Increasingly, quinolone resistance in foodborne pathogens has led to antibiotic treatment failure in gastrointestinal infection (Song et al., 2018).

*Salmonella* is an important foodborne zoonotic pathogen, associated with consumption of contaminated food (Akbar and Anal, 2015) and several studies have described transmission from livestock to humans (Bodhidatta et al., 2013; Fearnley et al., 2011; Prasertsee et al., 2019). Emerging AMR in NTS in food production is a serious issue for public health globally (Wang et al., 2019). Effective surveillance and molecular typing is necessary for source tracking of salmonellosis outbreaks (Revez et al., 2017). Pulsed field gel electrophoresis (PFGE), repetitive sequence based PCR (rep-PCR), and ribotyping have been used for *Salmonella* typing for several decades, but these classical serotyping methods cannot differentiate highly clonal *Salmonella* strains (Ranieri et al., 2013). Previously, we demonstrated that rep-PCR analysis can distinguish *Salmonella* serotypes, even when sampled from different sources (Prasertsee et al., 2019). Although this technique is appropriate for using for rapid identification of *Salmonella*, WGS offers increased resolution and more detail for molecular epidemiology studies (Prasertsee et al., 2016).

The Oxford Nanopore Technologies MinION long-read sequencer can provide rapid WGS for genomic epidemiology studies (Sang Chul et al., 2018). Our findings demonstrate high resolution for discriminating *Salmonella* clones, previously indistinguishable by rep-PCR analysis Both cg and wgMLST analysis clustered different *Salmonella* serotypes separately, with than 3% difference in loci within each serotype, reflecting high genetic relatedness of contemporaneous *Salmonella* isolates from meat and humans collected in the same geographical region. This is further evidence that *Salmonella* in meat products is a public health risk and an important source of human salmonellosis.

## 5. Conclusions

The MinION sequencing platform have advantages to perform the WGS in bacteria because of portability, affordability, real time base calling, and simplicity compared with other sequencing technologies. WGS data allows genome-wide genetic characterisation and high-resolution assessment of the relatedness of strains. Antimicrobial resistance genes can be identified and their location within the genome determined. Additioanlly, long-read sequencing allows reconstruction of plasmid sequence data. In this study, we investigated typed, putative *Salmonella* clones for improved core and whole-genome typing methods and characterisation of their AMR profiles. Our data reinforces that *Salmonella* isolates from meat can be transmitted to humans via the food chain. AMR genes were investigated and the resistant genotype correlated with displayed phenotypes. Several isolates were highly resistant and contained mobile plasmid elements that facilitaed the spread of AMR, including the last-line antibiotic, colistin.

## 6. Acknowledgements

This research was financially supported by the Royal Golden Jubilee (RGJ) Ph.D. Programme (PHD/0240/2558) and partially supported by Chiang Mai University. The authors would like to thank all scientists at Food Biotechnology Laboratory, National Science and Technology Development Agency (NSTDA) for their contribution. We would like to express many thanks to Dr.Supapon Cheevadhanarak, Dr.Sawannee Sutheeworapong, Dr.Songsak Wattanachaisereekul, and all staffs at Systems Biology and Bioinformatics research group (SBI) at King Mongkut’s University of Technology Thonburi (KMUTT) for excellent support and providing high performance computing servers.

